# Cyto/myeloarchitectural changes of cortical gray matter and superficial white matter in early neurodevelopment: Multimodal MRI study of preterm neonates

**DOI:** 10.1101/2021.03.16.435692

**Authors:** Shiyu Yuan, Mengting Liu, Sharon Kim, Jingda Yang, Anthony James Barkovich, Duan Xu, Hosung Kim

## Abstract

The developing cerebral cortex undergoes rapid microstructural and morphological changes throughout the third trimester. Recently, increased attention has been focused on the identification of imaging features that represent the underlying cortical cyto/myeloarchitecture driving intracortical myelination and the maturation of cortical gray matter (GM) and its adjacent superficial white matter (sWM). However, the characterization and spatiotemporal pattern of complex cyto/myeloarchitectural changes in this critical time period remain incompletely understood. Using 92 MRI scans from 78 preterm neonates (baseline: n□=□78, postmenstrual age=33.1±1.8 weeks; follow-up: n=14, 37.3±1.3), the current study leveraged combined T1/T2 intensity ratio and diffusion tensor imaging (DTI) measurements, including fractional anisotropy (FA) and mean diffusivity (MD), to characterize the cyto/myeloarchitectural architecture of cortical GM and its adjacent sWM in preterm neonates. DTI metrics during these weeks showed an overall linear developmental trajectory: FA decreased along with time in GM but increased in sWM; MD decreased in both GM and sWM. In contrast, T1/T2 measurements showed a distinctive parabolic developmental trajectory, revealing additional cyto/myeloarchitectural signature inferred. Furthermore, the spatiotemporal courses of T1/T2 ratio and DTI parameters were found to be regionally heterogeneous across the cerebral cortex, suggesting these imaging features’ specific relationship to regional cyto/myeloarchitectural maturation: faster T1/T2 ratio changes were found in the central, ventral, and temporal regions of GM and sWM, faster FA increases in anterior sWM areas, and faster MD decreases in GM and sWM central and cingulate areas. Taken together, our results may offer an explanation of the novel pattern of cyto/myeloarchitectural processes observed throughout the third trimester, including dendritic arborization, synaptogenesis, glial proliferation, as well as radial glial cell organization and apoptosis. Finally, T1/T2 ratio and DTI measurements were significantly associated with 1 year outcome scores of language and cognitive performance as well as perinatal clinical conditions, including intraventricular hemorrhage and chronic lung disease, demonstrating their potential as imaging biomarkers characterizing microstructural deviation in atypical neurodevelopment. Ultimately, with combined properties of cortical T1/T2 and DTI measurements, this study provides unique insights into the cellular processes and associated developmental mechanisms during the critical development of the third trimester.

## Introduction

Human brain maturation involves complex morphological changes of the cerebral cortex, including the expansion of white matter (WM) and gray matter (GM), thickness, surface area, and folding. Alterations of such cortical morphological properties, which can be characterized by magnetic resonance imaging (MRI), have been associated with neuropsychiatric [1, 2] and developmental disorders [3, 4] that originate in prenatal and postnatal periods. Such critical period of cortical development involves neuronal migration from the germinal matrix, rapid growth of pyramidal cells, synaptogenesis, and myelination, which establish cerebral connectivity and facilitate rapid synchronized communication among different brain regions [5].

Studies to date have demonstrated that brain myelination involves the transition from the pre-myelination to the myelination stage, when immature oligodendrocytes proliferate and initiate myelinogenesis through axon contact and wrapping [6]. These myelination events start during the late second or early third trimester in the brainstem and deeper brain structures, eventually progressing to more peripheral brain regions to include superficial cerebral white matter tracts. Alongside these myeloarchitectural changes include precisely tuned cytoarchitectural events, including neural apoptosis, radial glial proliferation, dendritic arborization, neurite (neurons, axons, and dendrites) outgrowth and lamination, and synaptogenesis [7, 8]. Recently, there has been increased spotlight on imaging features representing the underlying cortical cyto/myeloarchitecture that drives intracortical myelination and cortical maturation of GM and its adjacent superficial white matter (sWM) [9-17]. Indeed, disruptions of cyto/myeloarchitectural events may lead to pathological outcome and are important prognostic indicators of response to perinatal injuries [18, 19]. Impaired myelination and immature WM are also associated with abnormal neurodevelopment, preterm birth, and perinatal insults [20-22]. The precise temporospatial pattern of cortical cyto/myeloarchitectural events in cortical GM and its adjacent sWM throughout the critical time frame of the third trimester, however, remain unclear.

Myeloarchitectural patterns can be characterized *in vivo* using multicontrast T1-weighted (T1w) and T2-weighted (T2w) MRI, intensity values of which may vary along with changes of water molecules and tissue macromolecules composing brain myelin [5, 23, 24]. Use of the semi-quantitative ratio of T1w/T2w signal intensity enhances the sensitivity and contrast to myelin, while reducing inter-subject signal intensity bias [25] and has been helpful in studying brain connectivity [26] and cortical myelin maps [27, 28] in infants [29-31], healthy subjects [10, 11, 32], aging adults [33] and clinical populations [34-36]. Though, this imaging feature has yet to be employed in the analysis of preterm neonatal cortical development throughout the third trimester. Most developmental studies have only explored myeloarchitectural evolution from infancy to childhood and adulthood [37, 38].

The T1w/T2w intensity ratio, however, is not specific solely to myelination events and may reflect a combination of various cytoarchitectural changes, including associated with neurite (e.g., neurons and axons) proliferation and integrity. Thus, to enhance the specificity of information inferred, directional diffusivity measurements of diffusion tensor imaging (DTI) together with T1w/T2w signal intensities have the potential to distinguish among the variety of cyto/myeloarchitectural events. DTI generates metrics that are sensitive to the movement of water molecules, including fractional anisotropy (FA), mean diffusivity (MD), axial diffusivity (AD), and radial diffusivity (RD) (Table S1) [39-42].

While tract-based analyses of DTI have been investigated in preterm and term neonates, myelination and axonal development in cortical GM and sWM remain largely incompletely explored. A few studies have described serial quantitative measures with MRI in very preterm and healthy human infants [43-45]. However, these studies assessed myelination or axonal development visually [40] or quantitatively but in only several white and GM regions of interest, which were sampled using manually placed landmarks [39, 46]. A recent study explored cortical microstructural maturation in preterm neonates using FA and DTI mean kurtosis, but the measurements’ precise relation to timing and specificity of cyto-/myeloarchitectural events remain unclarified [47].

To attain a more complete picture of brain maturation, our study analyzed multimodal MRI features using combined features of T1w/T2w intensity ratio and DTI measurements. We aimed to characterize cortical cyto/myeloarchitecture of a preterm neonates scanned in the equivalent age of the third trimester. Firstly, we plotted the developmental trajectory of both cortical GM and sWM. Secondly, using a cortical region of interest (ROI) approach, we constructed differential spatiotemporal maps of T1/T2 and DTI to characterize regionally specific signatures of GM and sWM across the cerebral cortex. Thirdly, we examined the relationship between T1/T2 and DTI measures to better define the differential microstructural properties. And finally, we examined whether T1/T2 and DTI measures are associated with neurodevelopmental outcome and clinical conditions. We hypothesized that (1) the cyto/myeloarchitectural properties inferred by T1/T2 and DTI would be different in sWM and GM and (2) T1/T2 and DTI developmental trajectories are regionally distinct and indicative of spatiotemporally dependent developmental events of the cerebral cortex. With the goal of addressing the paucity of cyto/myeloarchiterature studies in the developing neonatal brain, our study importantly identifies microstructural changes that may provide a foundational understanding on neurodevelopment and clinical outcome.

## Materials and Methods

### Participants

Our dataset includes 78 preterm (mean postmenstrual age at birth [PMA]□=□28.23□±2.07 weeks; range 24–32 weeks), admitted to UCSF Benioff Children’s Hospital San Francisco between June 2011 and March 2019. Neonatal demographic and clinical information are provided in Table 1. All patients were scanned postnatally as soon as they were clinically stable (PMA at scan: 33.10 ± 1.88 weeks; range 28.80–35.70 weeks), and 39 patients were scanned again before discharge at late preterm age (PMA at scan: 37.31□±□1.70 weeks; range 33.14–41.98 weeks). However, many follow-up scans were excluded due to large motion artifact after quality control, resulting in a total of 92 T1w and T2w MRI scans, and DWI scans (78 baseline, 14 follow-up scans). Other exclusion criteria included 1) clinical evidence of a congenital malformation or syndrome, 2) congenital infection, and 3) newborns clinically unstable for transport to the MRI scanner. Parental consent was obtained for all cases following a protocol approved by the Institutional Committee on Human Research.

**Table 1.** Demographic and clinical characteristics for preterm neonates

| Demographic |  |
| --- | --- |
| Subjects (n) | 78 |
| Sex: males (n) | 39 |
| MRI scans (n) | 92 |
| GA at birth (weeks, mean ± SD) | 28.23 ± 2.07 |
| PMA at MRI |  |
| 1<sup>st</sup> scans | 33.10 ± 1.88 |
| 2<sup>nd</sup> scans | 37.31 ± 1.70 |

| Characteristic<sup>a</sup> | Number, (%) |
| --- | --- |
| **Maternal/Antenatal factors** |  |
| Maternal age, yrs | 31.56 ± 7.46 |
| Placenta previa | 4 (7.27) |
| Drug Abuse<sup>b</sup> | 11 (17.74) |
| Magnesium sulfate | 52 (77.61) |
| Exposure to prenatal steroids | 63 (88.73) |
| Chorioamnionitis | 11 (17.74) |
| **Delivery / Perinatal factors** |  |
| Twin | 30 (40.54) |
| Birth weight, gram | 1092.54 ± 355.94 |
| Caesarean section delivery | 50 (67.57) |
| **Postnatal factors** |  |
| Exposure to postnatal steroids | 7 (10.45) |
| Hypotension | 37 (53.62) |
| Infant infection<sup>c</sup> | 45 (62.50) |
| Patent ductus arteriosus (PDA)<sup>d</sup> | 30 (48.39) |
| Necrotizing enterocolitis<sup>e</sup> | 5 (7.25) |
| Duration of intubation, days | 9.12 ± 14.99 |
| Chronic lung disease | 17(24.64) |
| Seizures | 1(1.45) |
| Pneumothorax | 1(1.45) |

<sup>a</sup>Data presented as number (%), or mean ± standard deviation. <sup>b</sup>All subjects with maternal smoking (based on self-report) were exposed to marijuana, two were also exposed to tobacco. <sup>c</sup>Newborns with culture-positive sepsis, clinical signs of sepsis with negative blood culture, or meningitis were classified as having infection. <sup>d</sup>Newborns with clinical signs of PDA (prolonged systolic murmur, bounding pulses, and hyperdynamic precordium) and evidence of left-to-right flow through the PDA on echocardiogram were classified as having a PDA. <sup>e</sup>It was diagnosed based on Bell's stage II criteria [48]; *Numbers and percentages were derived from 78 preterm subjects whose data was recorded and known.* PMA: postmenstrual age.

### MRI and DTI acquisition

All participants underwent a brain MRI on a 3T GE Healthcare Discovery MR750 scanner. Infants were scanned under natural sleep after feeding. Sedation was used only with parental consent and when subjects initially moved to cause imaging artifact. Approximately 25% of subjects received sedation, usually pentobarbital in small doses. T1-weighted images were also acquired using sagittal 3D IR-SPGR (inversion time of 450 ms; FOV = 180 × 180 mm2; NEX = 1.00; FA = 15°), and were reformatted in the axial and coronal planes, yielding images with 0.7 × 0.7 × 1 mm^3^ spatial resolution. Diffusion weighted images were obtained using a single shot echo-planar sequence: 27 non-collinear diffusion gradients with a b-value of 1000 s/ mm^2^, three non-diffusion weighted (b0) volumes, echo/repetition time = 80/8000 ms, voxel size = 1.56 × 1.56 × 3 mm^3^, acquisition time = 4:08 min. An anatomical T2-weighted sequence (3D Cube) was also acquired: echo/repetition time = 65/2500 ms, voxel size = 0.63 × 1 × 0.63 mm^3^, acquisition time = 3 min 20 s.

### MRI-based diagnosis of neonatal brain injuries

A pediatric neuroradiologist (A.J.B.) blinded to patient history reviewed patient MRI scans including 3D T1- and axial T2-weighted sequences, as well as susceptibility weighted imaging (SWI) when available. Presence and severity of three leading drivers of neurodevelopmental deficits, i.e., intraventricular hemorrhage (IVH), ventriculomegaly (VM), and periventricular leukomalacia (PVL) or white matter injury (WMI), were visually scored (Table S2). Subsequently, IVH scores were binarized with “mild” representing grades 1–2, and “severe” representing grades 3–4; PVL/WMI and VM were categorized as “mild” for grade 1 and “severe” for grades 2–3. We merged infants with mild injuries and those with no injury into one none-mild injury group, since the mild injury and no injury groups exhibited no significant differences in the following analyses as well as previous studies [49, 50].

### Neurodevelopmental assessment

All infants were referred to the UCSF Intensive Care Nursery Follow-Up Program upon discharge for routine neurodevelopmental follow-up. Neurodevelopment was assessed using the Bayley-III, which was performed by unblinded pediatric psychiatrist at 12, 18, and 30 months’ corrected age. Follow-up was available in 26 of the 78 infants (33%).

### MR image processing of T1w and T2w imaging data

Three types of brain tissues (GM, WM, and CSF) were segmented using a joint label fusion method on T1w images. Next, the cortical surfaces of neonates were extracted from the segmentation using NEOCIVET pipeline [51, 52]. T2w images were then registered to the pre-processed T1w image using a rigid-body registration based on the similarity measure as mutual information. Then, T1/T2 ratio was computed using the manner introduced in a previous study [25] and sampled at vertices on the mid-cortical surface (=surface placed at the middle between GM/WM and GM/CSF borders) and the superficial white matter (sWM) surface (=surface placed at 2mm below the GM/WM border), which were triangulated with 81,924 vertices (163,840 polygons). These features were further re-sampled to the surface template using the transformation obtained in the surface registration, to allow inter-subject spatial correspondence.

### DTI processing

Motion artifacts and eddy current distortions in DTI data were first corrected by normalizing each directional volume to the b0 volume using FMRIB’s Linear Image Registration Tool (FLIRT) with 6° of freedom (DOF) [53]. The diffusion tensor was then calculated using a simple least square fit of the tensor model to the diffusion, and MD, AD, RD and FA maps were then generated.

To accurately estimate the intracranial correspondence between structural and diffusion weighted image spaces, a label-fusion approach [54] was used to mask out non-cortical structures for T1w and T2w images. The b0 images were initially skull-stripped using Brain Extraction Tool [55]. Given that regional intensity and geometric differences exist between T2w and b0 MR images [56], the ANTs software, a diffeomorphic mapping method with mutual information as the similarity measure, was used to warp the b0 images to T2 images. Based on the estimated transformation parameters in the rigid registration between T2w and T1w images, b0 volume (together with MD and FA volumes) was further transformed to T1 space for mapping the diffusion data on mid-cortical and sWM surfaces. Thus, FA, AD, RD and MD values were sampled on the mid-cortical and sWM surfaces for further inter-subject comparisons.

### Regional Analysis of T1/T2 ratio and DTI parameters

In this study, we use the surface-based parcellation, in which vertices that belong to the same anatomical structure are grouped. Here we used the automatic anatomical labeling (AAL) parcellation [57] mapped to the neonatal cortical template using the method described and evaluated in Kim et al. [51] to define cerebral cortical GM and sWM regions of interest (ROIs). After the surface registration, 81,924 vertices sampled on each individual surface were labeled into 76 ROIs. We then calculated the average of each of T1/T2 and DTI parameters sampled at vertices within each of 76 ROIs for each individual scan. These ROI mean features were used to analyze the pattern of spatiotemporal brain maturation from 28 to 42 GW.

### Spatiotemporally Varying Pattern of Developmental Trajectory

To evaluate the trajectory of cyto/myeloarchitectural maturation, linear mixed-effect regression models that addressed for repeated measurements (i.e., some subjects were scanned twice) were established for each T1/T2 ratio, MD, AD, RD or FA value as the dependent variable. In the models, we correlated each brain maturation metric with PMA at scan while including the following covariates as other independent variables: PMA at birth, and sex. We analyzed these regression models for the whole cortex and 76 cortical regions. Due to the nonlinear trend observed in T1/T2 values, we conducted a quadratic (=2^nd^ polynomial) regression model across the cohort, which resulted in smaller fitting errors than the linear model for this measurement.

### K-mean clustering to classify the cortical regions with respect to maturation rates and degrees

To facilitate the interpretation of the spatiotemporal maturation trajectory, we applied k-mean clustering to each imaging feature’s temporally changing map that was estimated using the aforementioned mixed-effect regression models. Using this unsupervised learning method, we classified the 76 cortical ROIs into two groups according to their two regional developmental characteristics: individual feature value at the term equivalent age of 40GW and the coefficient of feature value representing the maturation rate with respect to PMA across the cohort. Thus, the output of the k-mean clustering labeled all the 76 ROIs into either fast or slow maturation based on their maturation rate during the 3^rd^ trimester and the maturation level at term equivalent age.

### Effects of perinatal clinical factors on developmental trajectory

We aimed to assess effects of each of perinatal clinical factors on postnatal brain developmental trajectories. Each factor was dichotomized using the clinically defined categorization (Table S3). To assess the association of the given clinical variable with T1/T2, MD, AD, RD or FA values, we used a univariate mixed-effect linear model that addressed within-subject changes and inter-subject effects. We tested difference in each imaging feature values between the two groups dichotomized for each clinical variable while correcting for PMA.

### Prediction of neurodevelopmental outcome using machine learning and T1/T2 ratio and DTI parameters

To probe whether the cyto/myeloarchitectural maturation is associated with brain functional development, we examined whether T1/T2 and DTI parameters at neonatal scan could predict behavioral performance, i.e. the Bayley-III scores. For the prediction of Bayley scores, we employed random forest regression (RFR), a supervised machine learning algorithm, and obtained the ranking of the importance for each regional feature by using the out-of-bag samples [58]. Specifically, T1/T2, MD, AD, RD and FA values extracted from the mid surface and superficial WM surface from all 76 cortical regions were used as input features for the RFR model. To select the relevant features, we calculated the mean importance index per feature using the out-of-bag sampling method. We selected regional imaging features that showed the importance scores within the top 10%. We then trained the RFR model with the selected features to estimate Bayley scores. The prediction performance of the model was evaluated based on a leave-one-out cross-validation and the minimization of the objective function using Bayesian optimization. We empirically chose the following parameters that yielded the best performance: LSBoost as the ensemble-aggression method, number of learning cycles = 30, flag indicating to sample with replacement, and learning rate = 0.1.

## Results

### 1 Global trajectory of cyto/myeloarchitectural maturation (Figure.1)

For T1/T2 ratio developmental trajectory, the 2^nd^ order polynomial model showed a better fitting (mid-cortical surface: R = 0.443; superficial WM surface: R=0.580) than the linear model (mid: R = 0.378; superficial: R= 0.542). On the mid-cortical surface, the model showed that the global T1/T2 ratio declined along with PMA increase until around 31-32 gestational weeks (GW) and started increasing after 32 GW (fig.1). In the same fashion, the T1/T2 ratio sampled on the superficial WM (sWM) surface declined until around 31-32 GW and increased thereafter. For diffusion metrics, FA on the mid-cortical surface decreased linearly along with PMA increase (R=-0.504), whereas FA on sWM surface increased linearly (R=0.292). In addition, MD sampled on the mid-cortical surface presented a linearly decreasing trajectory along with PMA increase (R=-0.380). For the sWM surface, MD also showed a negative linear relationship with PMA (R=-0.559). Similar patterns were observed in RD and AD measurements (Supplementary Figure 1).

**Figure 1.**
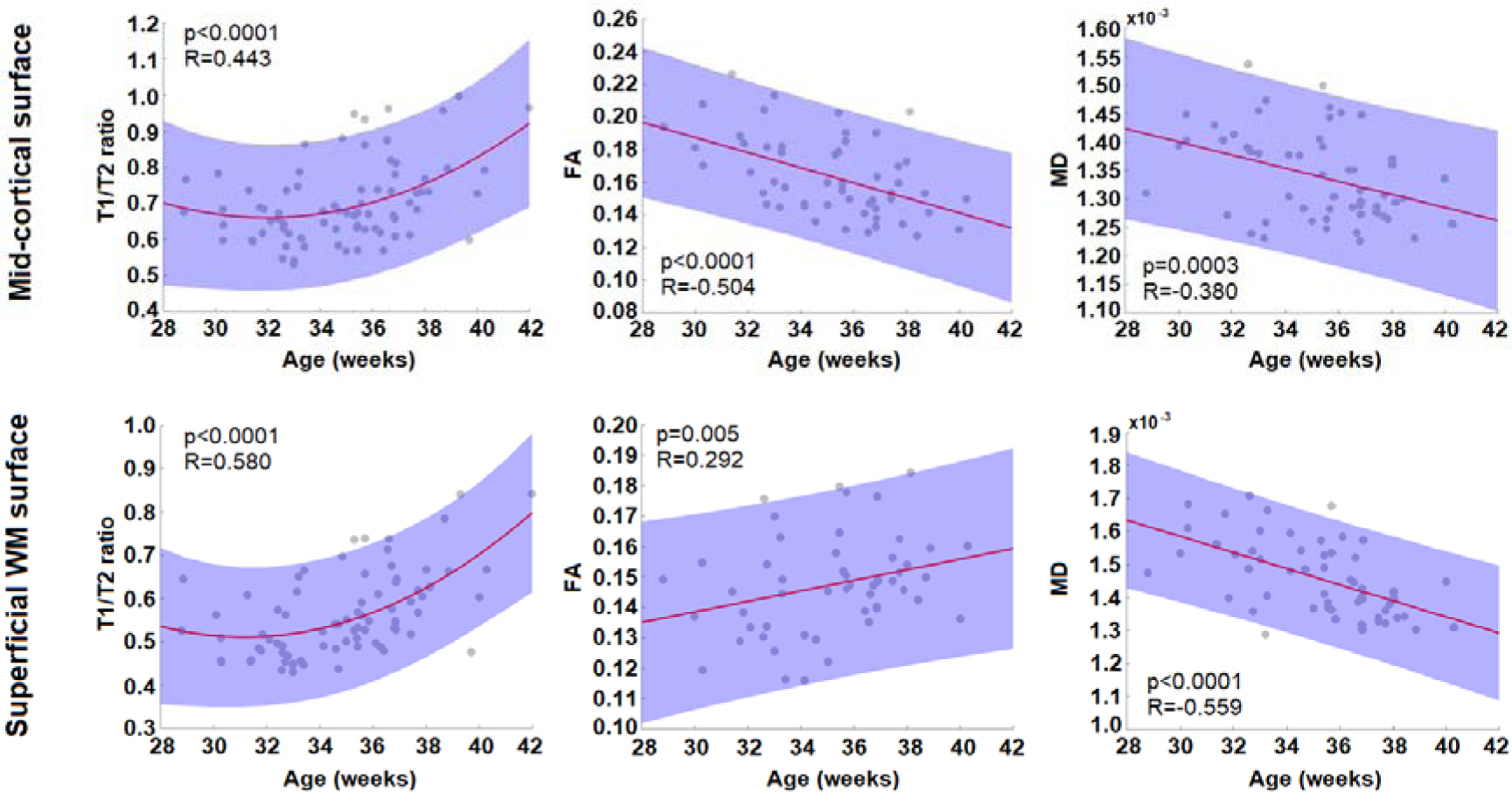
Global trajectory of cyto/myeloarchitectural maturation. While the trajectory of T1/T2 ratio metric was best fitted by parabolic (=2^nd^ order polynomial) curve in both cortical GM and sWM, that of DTI-FA and MD metrics was linear. In GM, we found a decrease of FA along with postmenstrual age (PMA) increase whereas an increase was found in sWM. The MD value linearly decreases over time in both GM and sWM.

### 2 Spatiotemporal Maturational Trajectory

Regional feature values of each T1/T2, FA, and MD represented using the fitted models are displayed at 28, 32, 36, and 40 GW in Figures 2-4, respectively. For each feature, we also included the results of the k-mean clustering that labeled all 76 ROIs into either fast or slow maturation.

**Figure 2.**
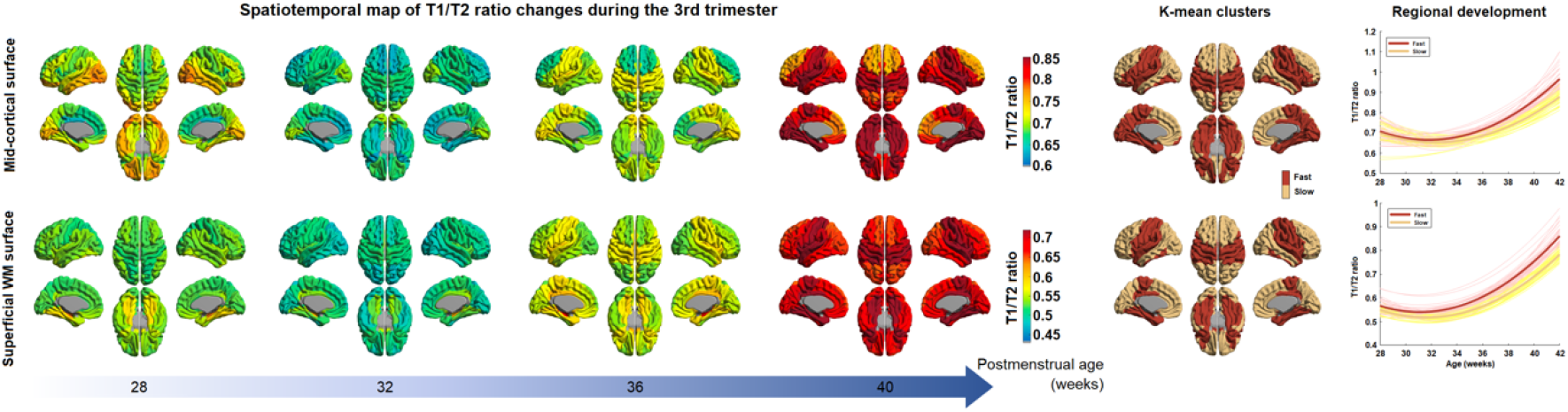
Spatiotemporal map of T1/T2 intensity ratio during the 3^rd^ trimester. The maps present T1/T2 values on the 2^nd^ polynomial curve fitted into the preterm data at each vertex. Mid-cortical surface (top) and superficial WM surface (bottom) are shown. The spatiotemporal pattern of T1/T2 intensity ratio is mapped in every 4 weeks from 28 to 40 week of PMA. The K-mean clustering results (the two most right columns) show regions with fast and slow maturation along with their developmental trajectories.

**Figure 3.**
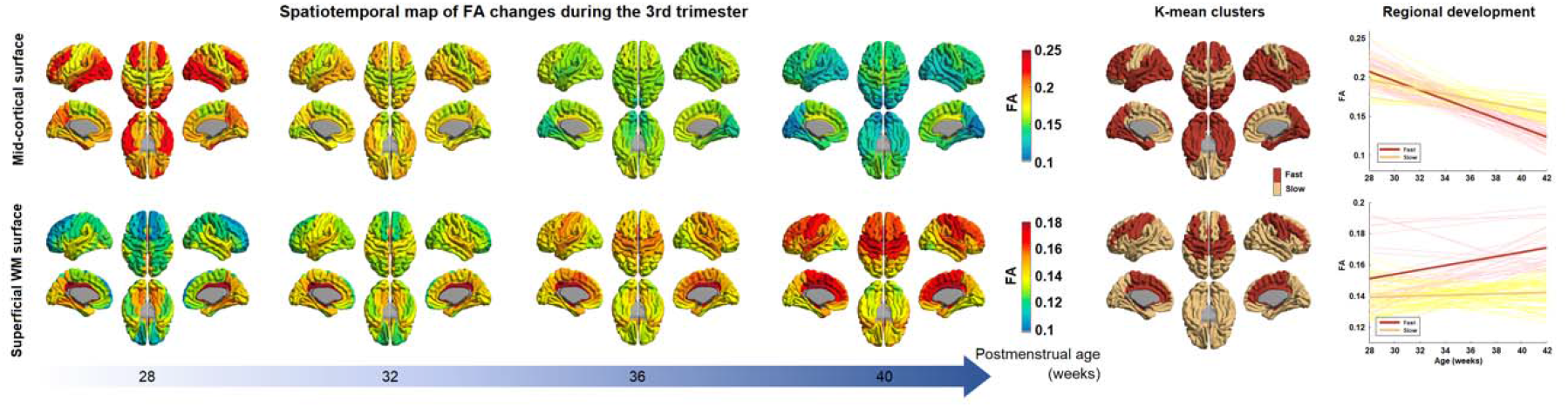
Spatiotemporal map of DTI fractional anisotropy (FA) during the 3^rd^ trimester. The maps present FA values on the linear model fitted into the preterm data at each vertex. Mid-cortical surface (top) and superficial WM surface (bottom) are shown. The spatiotemporal pattern of FA is mapped in every 4 weeks from 28 to 40 week of PMA. The K-mean clustering results (the two most right columns) show regions with fast and slow maturation along with their developmental trajectories.

**Figure 4.**
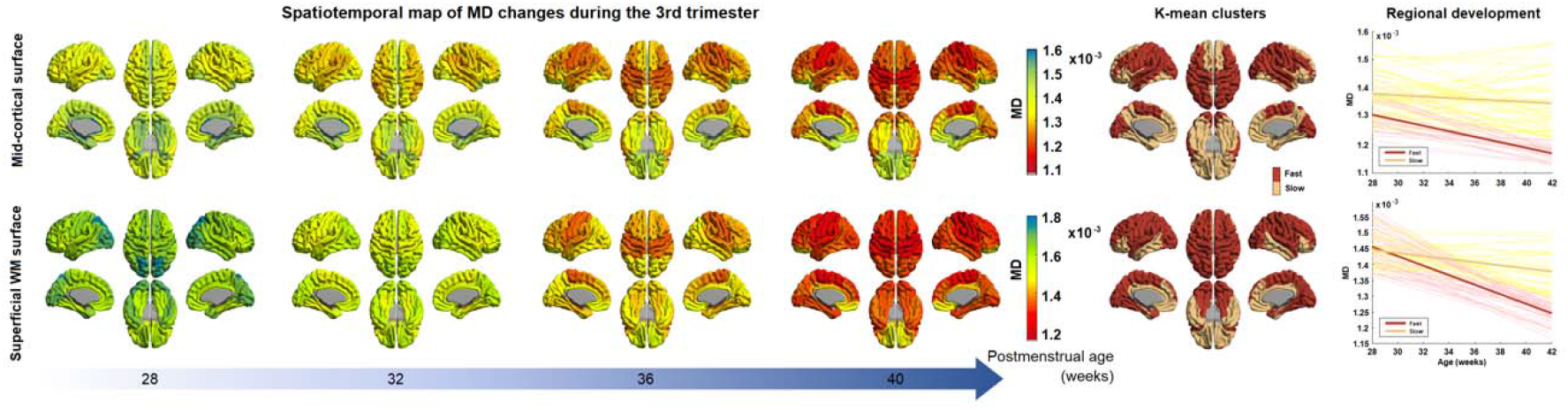
Spatiotemporal map of DTI mean diffusivity (MD) during the 3^rd^ trimester. The maps present MD values on the linear model fitted into the preterm data at each vertex. Mid-cortical surface (top) and superficial WM surface (bottom) are shown. The spatiotemporal pattern of MD is mapped in every 4 weeks from 28 to 40 week of PMA. The K-mean clustering results (the two most right columns) show regions with fast and slow maturation along with their developmental trajectories.

#### A T1/T2 ratio (**Figure 2**)

All cortical regions showed a similar “U-shape” parabolic developmental trajectory of T1/T2 values, with a significant association with PMA at scan (R>0.44; p<0.0001). The inflection point of T1/T2 values was found at around 32 GW for all regions consistently. The spatiotemporal pattern of T1/T2 changes on the mid-cortical surface across different PMA values is shown in Figure 2-top. At 28 GW, ventral regions, including lateral occipitotemporal and orbitofrontal, temporal and occipital cortices, showed relatively higher T1/T2 values than dorsal regions, including the superior frontal, central, and parietal lobe. By around 32 GW, T1/T2 values decreased to the inflection point in all regions. The occipital area exhibited a slightly higher T1/T2 ratio value than other regions. From 32 GW to 36 GW, T1/T2 ratio increases globally, especially in central and superior temporal cortices. At 40 GW, central and ventral cortical areas showed much higher T1/T2 values than other areas. Particularly, values in the medial occipital cortex (V1 and V2) were the highest among all ROIs. K-mean clustering showed that frontal, middle and inferior temporal, and lateral occipital cortices exhibit slow increases from 32 GW to 40 GW whereas superior, while basal and medial temporal, orbitofrontal, central and medial occipital cortical regions exhibit fast increases by 40 GW.

We also plotted the spatiotemporal pattern of T1/T2 changes occurring in the superficial WM over different PMA (Figure 2-bottom). At 28 GW, ventral regions, especially the medial occipitotemporal area and para-hippocampus, presented slightly higher T1/T2 values. By 32 GW, there were globally observed T1/T2 decreases to the lower inflection in sWM. At 36 GW, the T1/T2 ratio increased globally with more noticeable changes in the central, ventral, and temporal cortices. At 40 GW, all cortical regions displayed T1/T2 increases than 32 and 36 GW. K-mean clustering showed fast increases in lateral orbitofrontal, central, superior, basal and medial temporal regions.

#### B DTI-FA (**Figure 3**)

At 28 PMW, cortical FA values were higher in lateral frontal, lateral temporal and lateral occipital cortical regions relative to that in primary sensorimotor areas (Figure 3-top). However, we found the opposite pattern by 40 GW, in which lateral frontal, temporal and occipital regions showed rapidly decreased FA values by 40 GW, as also observed in K-mean clustering result. By contrast, the central, cingulate, medial frontal, and orbitofrontal regions underwent relatively slow FA decreases by 40 GW.

The sWM anisotropy underwent more heterogeneous and more dynamic changes compare to the anisotropy changes in GM (Figure 3-bottom). FA values increased with PMA in sWM, but different regions had distinct developmental patterns. The cingulate region maintains a relatively high FA value throughout different PMAs. The frontal lobe, especially the superior frontal cortex, has the lowest FA value at 28 GW, but it reached a relatively high FA value by 40 GW compare to other sWM brain regions. Generally, dorsal (superior) areas have lower FA values (<0.13) than ventral (inferior) areas in the initial stage, but dorsal areas reached higher FA values (>0.16) than ventral areas at 40GW, especially in central, superior, and medial frontal regions. Ventral areas underwent moderate anisotropy changes with increasing PMA. Noticeably, the precuneus, cuneus, parahippocampal and lateral occipitotemporal cortices displayed no FA changes or slight decreases as the PMA increased.

#### C DTI-MD (**Figure 4**)

In cortical GM (Figure 4-top), the lateral-dorsal cortical convexity, including lateral frontal, posterior cingulate, temporal and parietal cortical areas, maintained lower and more rapidly decreasing MD values than other cortical regions throughout development. By contrast, the medial-ventral cortical regions, including olfactory, basal/medial temporal, orbitofrontal and cingulate cortices, presented higher and more slowly decreasing MD values.

In sWM (Figure 4-bottom), the degree of MD at each cortical area differed from that seen in cortical GM. However, the overall spatiotemporal pattern of sWM was similar to the changes of cortical GM. One particular observation was that at 28 GW, the occipital region displayed relatively higher MD value than other regions. By 36 GW, central and cingulate WM regions showed a more rapid decrease in MD (=faster sWM development) than other regions. By 40 GW, most areas except the orbitofrontal, middle/inferior temporal, and anterior cingulate WMs (>0.0013) reached a similar low MD value (0.0011-0.0013).

### 3 Feature correlation

T1/T2 and DTI parameters may represent different aspects of cyto/myeloarchitectural maturation (Van Essen et al., 2011) while any one of these parameters may not fully explain all aspects of cortical maturation. Particularly, the T1/T2 ratio that has been used to explain the degree of cortical myelination in adult brains may also be affected by cytoarchitectural changes coinciding in the third trimester and early postnatal period, including selective elimination of neuronal processes through cell death, and proliferation and differentiation of glia[7]. To facilitate the relationship between T1/T2 and DTI parameters, we computed the correlation between T1/T2 and DTI measurements for each cortical ROI (38 ROIs after merge the left and right hemispheres; Figure 5).

**Figure 5.**
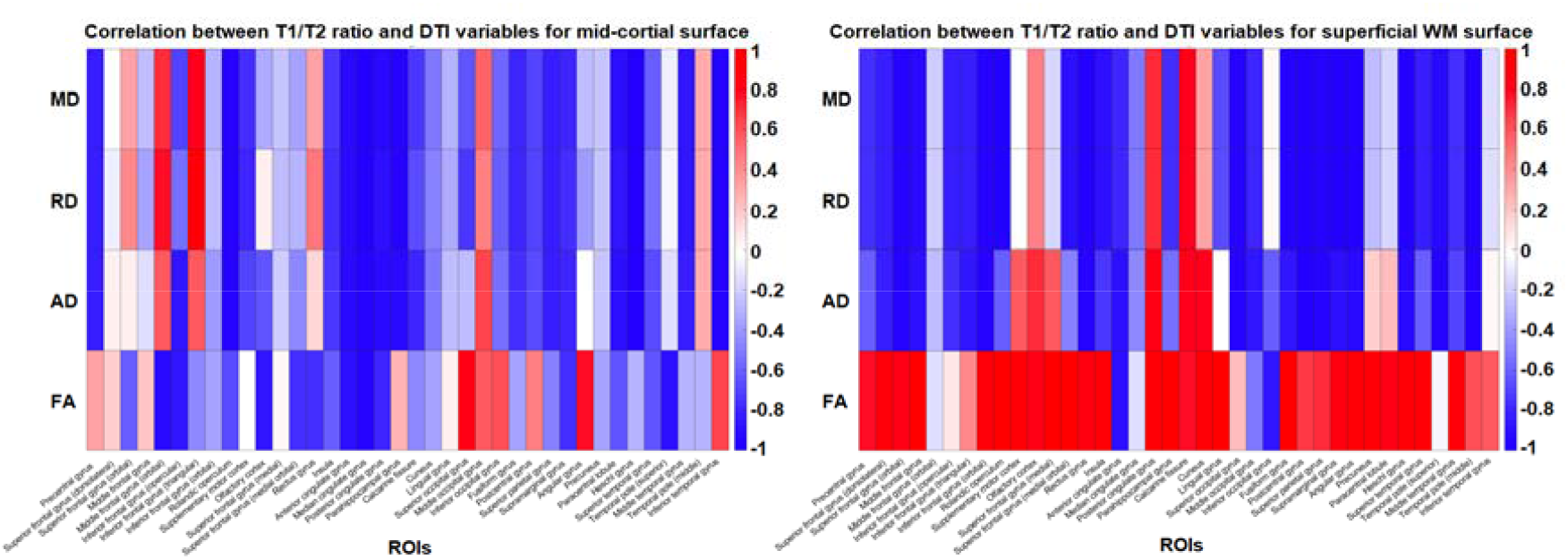
Relationship between T1/T2 intensity ratio and DTI-parameters. The color represent either negative (blue) or positive (red) correlation between T1/T2 intensity ratio and each of MD, RD, AD and FA.

In cortical GM, T1/T2 ratio showed similar patterns in regional correlations with DTI-MD, RD and AD, in which about half of cortical regions displayed negative correlations (17 ROIs (44%) with a large effect size of correlation coefficient |r|=0.5 or higher) and only few showed positive correlations (3 ROIs (7%) with r>0.05). By contrast, T1/T2 ratio showed a mixture of positive and negative correlations with FA (7 ROIs (18%) with r>0.5; 15 ROIs (39%) with r<-0.5. For example, T1/T2 ratio in precentral, parahippocampal, angular, and inferior temporal regions, diffusivity correlated positively with FA but negatively with diffusivity. Contrarily, T1/T2 values in orbital, inferior frontal, superior occipital, temporal pole regions correlated negatively with FA but positively with diffusivity metrics. In superior temporal, cingulate, supramarginal gyrus, central lobular, precuneus and cuneus regions, T1/T2 ratio correlated negatively with both diffusivity metrics and FA. In the middle occipital gyrus, T1/T2 ratio correlated positively with both diffusivity metrics and FA.

In sWM, T1/T2 ratio negatively correlated with diffusivity metrics (MD, AD, RD) in the majority of regions (27 ROIs (71%) with r<-0.5). Only the posterior cingulate, cuneus and olfactory cortices showed significant positive correlations between T1/T2 and diffusivity metrics (3 ROIs (7%) with r>0.5). By contrast, T1/T2 ratio positively correlated with FA in most regions (30 ROIs (79%) with r>0.5. The lateral occipital, cingulate, orbitofrontal cortices showed negative correlations with FA (3 ROIs (7%) with r<-0.5).

### 4 The association between clinical factors and imaging features of cortical development

Our univariate analyses showed that severe brain injury (intraventricular hemorrhage IVH) was related to increased diffusivity in both cortical GM (parahippocampal cortex) and sWM (parahippocampal, and lingual cortices) because the severely injured group presented a higher MD value than the unaffected group (Figure 6). The sWM showed more pronounced and extended damage in relation to severe brain injury compared to GM. In cortical GM, higher FA was significantly associated with the occurrence of chronic lung disease (CLD) (Figure 6). This significance was not observed in sWM. CLD affected lateral/medial prefrontal cortices and anterior temporal and mesiotemporal cortices.

**Figure 6.**
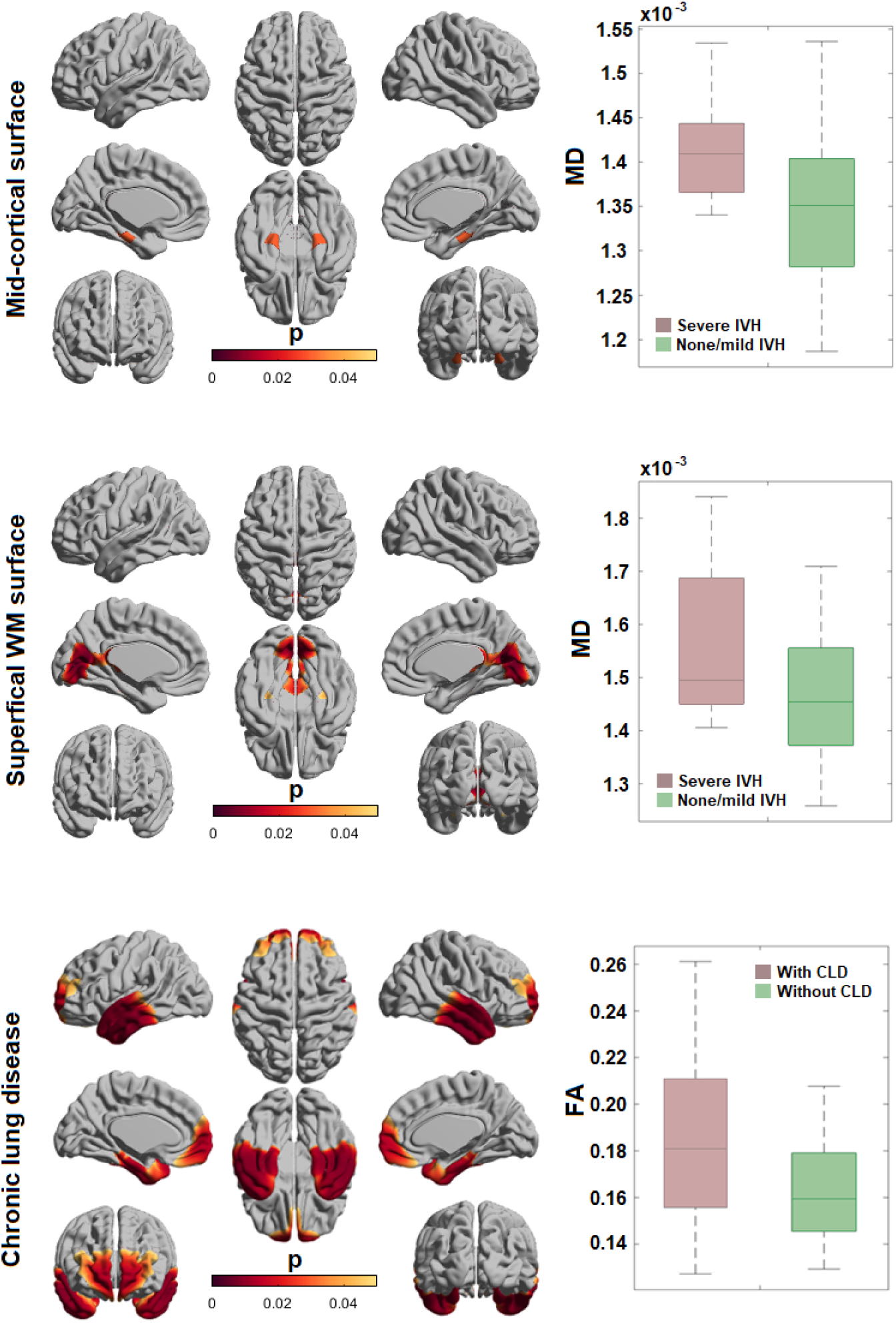
The effect of perinatal brain injury and postnatal clinical factors on cortical GM and superficial WM maturation in preterm neonates. The presence of severe intraventricular hemorrhage is associated with higher mean diffusivity in GM and WM regions. Postnatal chronic lung disease are associated with higher fractional anisotropy in cortical GM regions.

### 5 Cognitive score prediction

The multivariate random forest regression (RFR) analysis with feature selection and leave-one-out cross-validation (Figure 7) showed that the RFR models, when using the most important predictive features, accurately predicted neurodevelopmental cognitive (r=0.48; p=0.011) and language (r=0.44; p=0.03) scores measured at 12 months. The RFR model did not significantly predict motor scores at 12 months (p>0.1). Cortical features that were identified as important predictors consistently in all leave-one-out cycles were mapped in Figure 7B. Note that diffusivity features (MD, AD, RD) were not included in the significant predictors of neurodevelopmental outcomes. Important features predicting cognitive performance at 12 months largely overlapped with those predicting language function. This is possibly because subjects showed a significant correlation between their cognitive and language functional scores (r=0.63), and our RFR models that were separately applied to the prediction of each function did not account for the covariance of these two functions.

**Figure 7.**
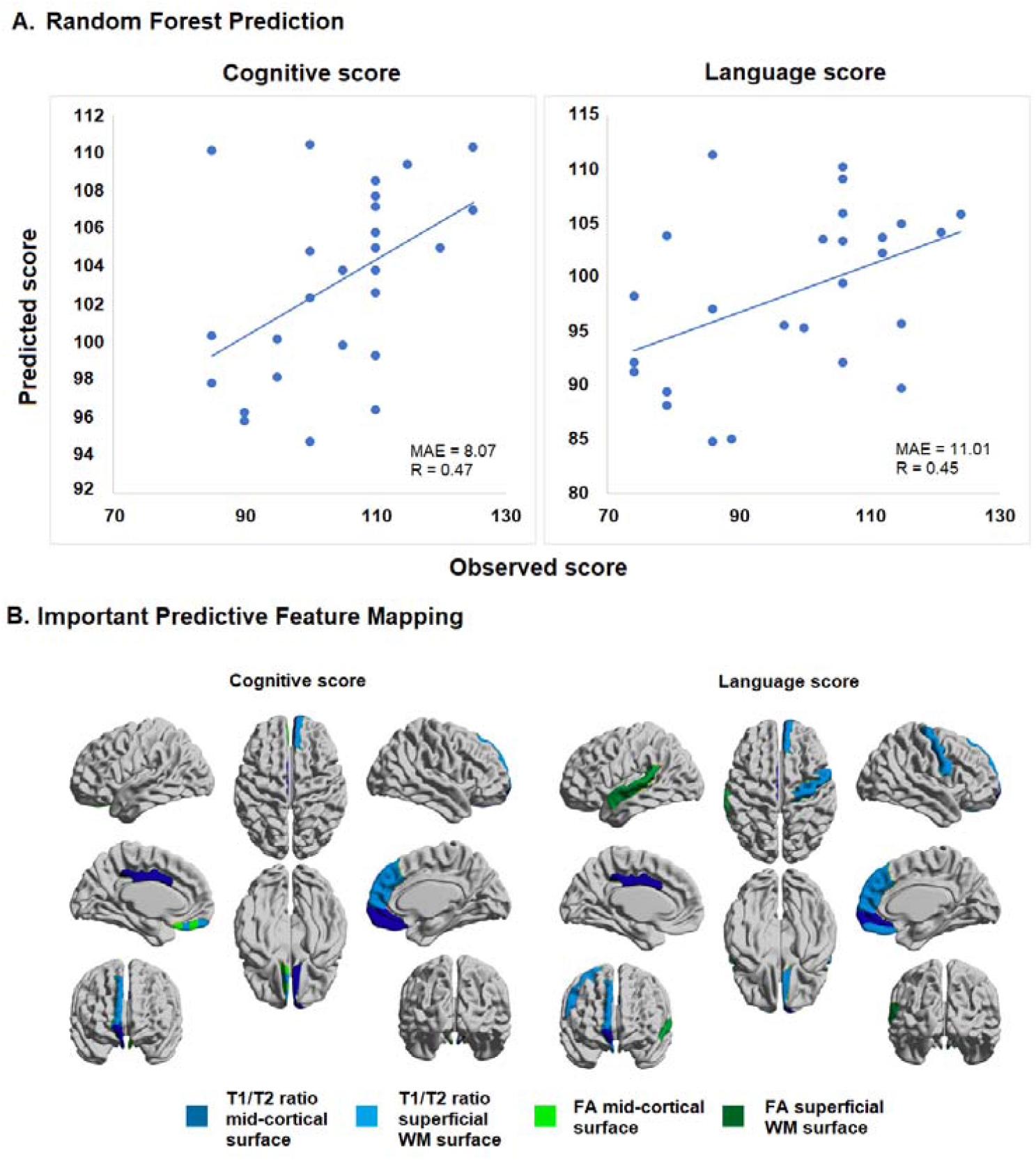
A) Random forest regression (RFR) significantly predicted neurodevelopmental assessment scores for cognitive (r=0.48; p=0.011, mean absolute error [MAE]: 8.07) and language (r=0.44; p=0.03, MAE=11.01) scores measured at 12 months. B) Important features predicting cognitive and language performance at 12 months were mapped onto cortical surfaces.

The mid-cortical and sWM imaging features did not predict neurodevelopmental outcomes measured at 18 and 30 months.

## Discussion

### Summary of Findings

Our study revealed differential developmental trajectories of global T1/T2 as well as DTI FA and MD across the cerebral cortex in both cortical GM and sWM, indicating that the microstructural properties inferred by the two measurements are distinct (Figure 1). Furthermore, the microstructural developmental courses of T1/T2 and DTI parameters were found to be regionally varying across the cerebral cortex, suggesting their sensitivity and specificity to important local cyto/myeloarchitectural changes (Figure 2-5). Finally, T1/T2 and DTI-FA were significantly associated with neurodevelopmental outcomes and clinical conditions, demonstrating their potential in quantifying aberrant neurodevelopment (Figure 6-7).

### Developmental Trajectory in Cortical Gray Matter and Associated Cyto/Myeloarchitectural Processes

Our results showed that global FA values decrease in cortical GM throughout the third trimester (Figure 1). Decreasing global FA in cortical GM is largely consistent with previous findings [46, 59] and has been interpreted as due to the lack of diffusion barriers upon radial glia apoptosis [47]. The higher anisotropy in early developing cortical GM may also be due to the predominance of radial glial fibers (RGF), or neurons with simple axons and an underdeveloped dendritic tree, while the decline over the period from 26 to 40 weeks may reflect the expansion of dendritic tree and axonal ramifications alongside the regression of RGFs. Accordingly, one recent study using DTI metrics proposed a hypothetical model, in which cortical FA values are predominantly influenced by RGF apoptosis [47].

In contrast to FA trends, we found a distinct parabolic trend of T1/T2 values in cortical GM (Figure 1), which may capture more subtle cyto/myeloarchitectural changes. Specifically, we found that T1/T2 values decrease from 28-32 weeks but increase parabolically from 32-40 weeks. The initial T1/T2 decrease between 28-32 weeks may be principally driven by radial glial cell organization whereas cyto/myeloarchitectural events may predominate to drive T1/T2 increases between 32-40 weeks as the radial cells undergo apoptosis and differentiate into neuronal precursors. In fact, several studies have already proposed that T1/T2 is not only sensitive to intracortical myelination properties [11, 25] but also to cortical dendrite pathology [60-62]. Thus, T1/T2 increases may reflect a variety of cyto/myeloarchitectural changes, including the elaboration of dendritic and axonal ramifications, establishment of synaptic contacts, selective elimination of neuronal processes through cell death, and proliferation and differentiation of glia [7].

### Developmental Trajectory in Adjacent Superficial White Matter and Associated Cyto/Myeloarchitectural Processes

Our findings revealed that the developmental trajectory of FA values in sWM is different from that of cortical GM, indicating differential developmental roles of the two tissue subtypes despite their close proximity. Specifically and consistently with prior studies [46, 63], we found that FA values increased in sWM. In contrast to deep WM that plays an important role in subcortical and remote cortical connectivity, sWM is strongly influenced by cortical GM, particularly short cortical association fibers (also called arcuate or “U”-fibers), which connect adjacent gyri and comprise the majority of corticocortical WM connections [15]. FA increases in sWM might thus indicate stronger influence from axonal packing organization and myelination on short corticocortical WM connections.

This is further supported by our findings on a less parabolic and more linear trajectory of T1/T2 in sWM, implicating the expansion of myeloarchitectural events. Throughout development, myelination events occur through the oligodendroglial lineage that begins with the oligodendroglial progenitor and continues successively with the preoligodendrocyte, the immature oligodendrocyte, and the mature oligodendrocyte. In early third trimester, preoligodendrocyte differentiation into immature oligodendrocyte predominate. And during the later stage of the third trimester, axonal ensheathment by premyelinating oligodendrocytes predominate [6, 64] contributing to premyelination anisotropy [65, 66] and increases in T1/T2 intensity [25].

Our exploration of the relationship between T1/T2 and FA (Figure 5) showed that, while T1/T2 values were mostly negatively correlated with FA in cortical GM, they were significantly positively correlated with FA in sWM. This supports our aforementioned hypothesis that information inferred by T1/T2 values in cortical GM is different from that inferred by FA. More specifically, a negative correlation between T1/T2 and FA in cortical GM regions supports our hypothesis in which cerebral cortical GM undergoes combined cyto/myeloproliferative events beyond simplified events related to radial glial cell apoptosis. By contrast, a positive correlation between T1/T2 and FA in sWM confirms that information inferred by both FA and T1/T2 may overlap in sWM i.e., myeloproliferative events.

### Consistent Decreases in MD for Cortical Gray Matter and Adjacent Superficial White Matter

Our findings demonstrate that global MD values decrease in both cortical GM and sWM throughout the third trimester. Several developmental studies have found similar trends in WM where decreasing MD is coupled with a rise in FA in preterm infants [67-71], term infants [72, 73], and adolescence [74, 75]. In an intriguing contrast, cortical GM exhibits overall decreases in both MD and FA, as shown in prior studies on preterm subjects [46, 59]. Decreasing MD in GM can be reflective of more complex microstructural barriers to diffusion due to cyto/myeloproliferative events. These changes may be explained by decreasing diffusivity of the radial direction (radial diffusivity, RD; Figure S1), implicating stronger influence of myelination events in this process. Indeed, it is well-established that myelination events progress throughout the third trimester, adolescence, and up to adulthood where these changes plateau [76].

### Heterogeneous Spatiotemporal Patterns of Cortical and sWM T1/T2 and DTI-FA and MD Measurements

Our results suggest that T1/T2 and DTI-FA and MD developmental trajectories are unique and differential across cortical regions. Such heterogenous developmental profiles possibly lead to unique cortical regions important for distinct neural circuitry and functional systems [77-79]. Among these patterns, one noteworthy finding is that FA decreased more rapidly in the frontal, temporal, and occipital cortical GM areas by the 40th week of gestation, indicating that these regions underwent more intense cytoarchitectural changes throughout development. By contrast, slower FA decreases in the central, cingulate, and medial frontal regions suggest that they undergo maturation even earlier. Relatively low FA values at 30 weeks with slower decrease thereafter in the primary sensorimotor cortex also suggest its earlier development compared to higher-order prefrontal cortical regions, as the folding of sensorimotor cortices is most active during the late 2^nd^ and early 3^rd^ trimester before 30 weeks [51, 80]. Altogether, these regionally specific changes demonstrate a developmental gradient moving along a rostral and caudal direction to the frontal and occipital lobe, which is consistent with patterns documented in prior histopathological findings [76] as well as the heterogeneity of FA decreases observed in prior DTI studies [81-83].

Spatiotemporally, MD values decreased the slowest in the lateral cortical convexity of both sWM and GM, including lateral frontal, temporal, and parietal regions, indicating their faster maturation of cortical premyelinated axons relative to other regions. Paralleling what has been observed in prior studies [84, 85], these regional variations in the timing of this developmental trajectory follow a roughly posterior-to-anterior trend.

### Cortical T1/T2 and DTI Measurements Associated with Neurodevelopmental Outcome and Clinical Characteristics

Importantly, our study also found that T1/T2 and FA accurately predicts neurodevelopmental assessment scores for cognitive and language function measured at 12 months. MD was not a significant predictor for outcomes in preterm neonates potentially due to its informational redundancy as it positively and strongly correlated with T1/T2. Brain regions that accurately predicted cognitive scores included the medial frontal cortex and cingulate, which are important for learning, decision-making, and memory. Language scores were also predicted by the corresponding region of the superior temporal lobe, which is involved in the comprehension of language. Interestingly, features that predicted cognitive performance at 12 months largely overlapped with those predicting language function, indicating the widespread involvement of cortical regions in establishing synaptic architecture for multiple domains in early neurodevelopment. Our findings on the significant correlation between cognitive and language scores at 12 months also support the notion that multiple cortical regions may synchronize into a network of domains, as prior findings have indicated the formation of functional and structural networks throughout preterm development [86]. However, our models that were separately applied to the prediction of each of cognitive and language function did not account for the covariance of these two functions, limiting the specificity of imaging features to different functional domains. Importantly, the significant prediction of language and cognitive scores using neonatal cyto/myeloarchitecutural features was only found in the early 12 months of infancy but not for later years i.e., 24-30 months. This implicates that impairments in cyto/myeloarchitectural maturations may recover as these processes continue throughout adolescence and thus may not be as sensitive to long-term neurodevelopmental outcomes. By contrast, other features of neonatal brain MRI such as connectivity or morphological alterations i.e., cortical folding may be more permanent and significantly associated with long-term impairments as supported in previous studies [87, 88].

Additionally, our analyses demonstrated that severe IVH is associated with increased diffusivity in both cortical GM (parahippocampal cortex) and sWM (parahippocampal, and lingual cortices) (Figure 6), suggesting that IVH may impede regionally specific microstructural maturation. These findings align with prior studies suggesting that medical risk factors, including IVH, may be an important predictor of poor neurologic outcome [89-91]. Our study also demonstrated that, in cortical GM, higher FA values were significantly associated with the occurrence of CLD (Figure 6) i.e., lateral/medial prefrontal cortices, anterior temporal and mesiotemporal cortices. CLD occurs in preterm infants who are exposed to prolonged mechanical ventilation and oxygen therapy for pulmonary complications that can lead to hypoxia-ischemia, inflammation, germinal matrix injury, diffuse WM injury, and diffuse GM injury [92, 93]. These complications have been shown to be associated with poor neurodevelopmental outcomes, including motor [94], cognitive [95], and language deficits [96-98].

### Limitations and Future Directions

Our study has several limitations. Specifically, our ability to fully characterize cortical cyto/myeloarchitectural development was limited by a small sample size of 78 neonates, in which not every subject had a successful follow-up scan. Thus, this study is limited by its partial longitudinal design despite its mixed-effect models for statistical analyses. Additionally, this study analyzed only preterm neonates, which may have impacted our ability to assess normal development, as preterm birth itself may alter brain development even after correcting for postmenstrual age at birth. Due to a small sample size, the effects of the related risk factors, including variables listed in Table 1, were analyzed only in a univariate fashion without adjusting for their combined effects. Nevertheless, our study contributes to the growing body of research that leverages MRI to examine preterm cortical development throughout the third trimester [59, 83, 99].

Finally, although our innovative approach in combining T1/T2 and DTI parameters effectively characterized the dynamics of cortical development, the proposed cyto/myeloarchitectural models and its clinical utility require future studies to validate with neurohistological measurements, including neuronal density, glial proliferation, or dendritic arborization.

## Conclusion

Overall, with noninvasive properties of cortical T1/T2 and DTI-FA and MD measurements, this study provides an innovative approach to the longitudinal spatiotemporal mapping of cortical cyto/myeloarchitecture, offering unique insights into the cellular processes and associated developmental mechanisms during the third trimester. T1/T2 measurements complement DTI-derived FA measurements through its unique parabolic trend encompassing the timing of cyto/myeloarchitectural events, including radial glial cell organization and apoptosis, dendritic arborization, myelination, synaptic formation, and potentially brain circuit emergence. While the exact neuroanatomical underpinning of cortical T1/T2 and DTI metrics in cortical GM and its adjacent sWM is not completely known, the unique combination of DTI linear trajectories and T1/T2 parabolic trajectories contribute new insight on cyto/myeloarchitectural events in brain development. And finally, our study demonstrates that DTI and T1/T2 measurements are sensitive to impacts from clinically adverse events and may serve as clinical biomarkers of neurodevelopmental outcome at early infancy.

## Acknowledgements

This study was supported by the National Institutes of Health grants (P50NS035902, P01NS082330, R01NS046432, R01HD072074; P41EB015922; U54EB020406; U19AG024904; U01NS086090; 003585-00001) and BrightFocus Foundation Award (A2019052S).

## References

1. Li, G., et al., Cortical thickness and surface area in neonates at high risk for schizophrenia. Brain Struct Funct, 2016. 221(1): p. 447–61.

2. Sierra, M., et al., A structural MRI study of cortical thickness in depersonalisation disorder. Psychiatry Res, 2014. 224(1): p. 1–7.

3. Ecker, C., et al., Is there a common underlying mechanism for age-related decline in cortical thickness? Neuroreport, 2009. 20(13): p. 1155–60.

4. Pacheco, J., et al., Greater cortical thinning in normal older adults predicts later cognitive impairment. Neurobiol Aging, 2015. 36(2): p. 903–8.

5. Laule, C., et al., Magnetic resonance imaging of myelin. Neurotherapeutics, 2007. 4(3): p. 460–84.

6. Back, S.A., et al., Late oligodendrocyte progenitors coincide with the developmental window of vulnerability for human perinatal white matter injury. J Neurosci, 2001. 21(4): p. 1302–12.

7. Mrzljak, L., et al., Prenatal development of neurons in the human prefrontal cortex: I. A qualitative Golgi study. J Comp Neurol, 1988. 271(3): p. 355–86.

8. Rakic, P., Developmental and evolutionary adaptations of cortical radial glia. Cereb Cortex, 2003. 13(6): p. 541–9.

9. Uddin, M.N., et al., Comparisons between multi-component myelin water fraction, T1w/T2w ratio, and diffusion tensor imaging measures in healthy human brain structures. Sci Rep, 2019. 9(1): p. 2500.

10. Grydeland, H., et al., Intracortical myelin links with performance variability across the human lifespan: results from T1- and T2-weighted MRI myelin mapping and diffusion tensor imaging. J Neurosci, 2013. 33(47): p. 18618–30.

11. Shafee, R., R.L. Buckner, and B. Fischl, Gray matter myelination of 1555 human brains using partial volume corrected MRI images. Neuroimage, 2015. 105: p. 473–85.

12. Norbom, L.B., et al., Maturation of cortical microstructure and cognitive development in childhood and adolescence: A T1w/T2w ratio MRI study. Hum Brain Mapp, 2020. 41(16): p. 4676–4690.

13. Wu, M., et al., Development of superficial white matter and its structural interplay with cortical gray matter in children and adolescents. Hum Brain Mapp, 2014. 35(6): p. 2806–16.

14. Oyefiade, A.A., et al., Development of short-range white matter in healthy children and adolescents. Hum Brain Mapp, 2018. 39(1): p. 204–217.

15. Kirilina, E., et al., Superficial white matter imaging: Contrast mechanisms and whole-brain in vivo mapping. Sci Adv, 2020. 6(41).

16. Phillips, O.R., et al., Superficial white matter: effects of age, sex, and hemisphere. Brain Connect, 2013. 3(2): p. 146–59.

17. Timmler, S. and M. Simons, Grey matter myelination. Glia, 2019. 67(11): p. 2063–2070.

18. Hinojosa-Rodriguez, M., et al., Clinical neuroimaging in the preterm infant: Diagnosis and prognosis. Neuroimage Clin, 2017. 16: p. 355–368.

19. Volpe, J.J., Brain injury in premature infants: a complex amalgam of destructive and developmental disturbances. Lancet Neurol, 2009. 8(1): p. 110–24.

20. Volpe, J.J., et al., Reprint of “The developing oligodendrocyte: key cellular target in brain injury in the premature infant”. Int J Dev Neurosci, 2011. 29(6): p. 565–82.

21. Ortinau, C. and J. Neil, The neuroanatomy of prematurity: normal brain development and the impact of preterm birth. Clin Anat, 2015. 28(2): p. 168–83.

22. Inder, T.E., et al., Abnormal cerebral structure is present at term in premature infants. Pediatrics, 2005. 115(2): p. 286–94.

23. Clark, V.P., E. Courchesne, and M. Grafe, In vivo myeloarchitectonic analysis of human striate and extrastriate cortex using magnetic resonance imaging. Cereb Cortex, 1992. 2(5): p. 417–24.

24. Barkovich, A.J., Magnetic resonance techniques in the assessment of myelin and myelination. J Inherit Metab Dis, 2005. 28(3): p. 311–43.

25. Glasser, M.F. and D.C. Van Essen, Mapping human cortical areas in vivo based on myelin content as revealed by T1- and T2-weighted MRI. J Neurosci, 2011. 31(32): p. 11597–616.

26. Ma, Z. and N. Zhang, Cross-population myelination covariance of human cerebral cortex. Hum Brain Mapp, 2017. 38(9): p. 4730–4743.

27. Glasser, M.F., et al., Trends and properties of human cerebral cortex: correlations with cortical myelin content. Neuroimage, 2014. 93 Pt 2: p. 165–75.

28. Van Essen, D.C., et al., The WU-Minn Human Connectome Project: an overview. Neuroimage, 2013. 80: p. 62–79.

29. Chen, H., et al., White Matter Fiber-based Analysis of T1w/T2w Ratio Map. Proc SPIE Int Soc Opt Eng, 2017. 10133.

30. Lee, K., et al., Early Postnatal Myelin Content Estimate of White Matter via T1w/T2w Ratio. Proc SPIE Int Soc Opt Eng, 2015. 9417.

31. Soun, J.E., et al., Evaluation of neonatal brain myelination using the T1- and T2-weighted MRI ratio. J Magn Reson Imaging, 2017. 46(3): p. 690–696.

32. Chopra, S., et al., More highly myelinated white matter tracts are associated with faster processing speed in healthy adults. Neuroimage, 2018. 171: p. 332–340.

33. Teubner-Rhodes, S., et al., Aging-Resilient Associations between the Arcuate Fasciculus and Vocabulary Knowledge: Microstructure or Morphology? J Neurosci, 2016. 36(27): p. 7210–22.

34. Iwatani, J., et al., Use of T1-weighted/T2-weighted magnetic resonance ratio images to elucidate changes in the schizophrenic brain. Brain Behav, 2015. 5(10): p. e00399.

35. Chao, L.L., et al., Preliminary Evidence of Increased Hippocampal Myelin Content in Veterans with Posttraumatic Stress Disorder. Front Behav Neurosci, 2015. 9: p. 333.

36. Nakamura, K., et al., T1-/T2-weighted ratio differs in demyelinated cortex in multiple sclerosis. Ann Neurol, 2017. 82(4): p. 635–639.

37. Brody, B.A., et al., Sequence of central nervous system myelination in human infancy. I. An autopsy study of myelination. J Neuropathol Exp Neurol, 1987. 46(3): p. 283–301.

38. Deoni, S.C., et al., Cortical maturation and myelination in healthy toddlers and young children. Neuroimage, 2015. 115: p. 147–61.

39. Anjari, M., et al., Diffusion tensor imaging with tract-based spatial statistics reveals local white matter abnormalities in preterm infants. Neuroimage, 2007. 35(3): p. 1021–7.

40. Counsell, S.J., et al., Specific relations between neurodevelopmental abilities and white matter microstructure in children born preterm. Brain, 2008. 131(Pt 12): p. 3201–8.

41. Duerden, E.G., et al., Tract-Based Spatial Statistics in Preterm-Born Neonates Predicts Cognitive and Motor Outcomes at 18 Months. AJNR Am J Neuroradiol, 2015. 36(8): p. 1565–71.

42. Beaulieu, C., The Biological Basis of Diffusion Anisotropy, in Diffusion MRI. 2014, Academic Press. p. 155–183.

43. Schneider, J., et al., Evolution of T1 Relaxation, ADC, and Fractional Anisotropy during Early Brain Maturation: A Serial Imaging Study on Preterm Infants. AJNR Am J Neuroradiol, 2016. 37(1): p. 155–62.

44. Counsell, S.J., et al., MR imaging assessment of myelination in the very preterm brain. American Journal of Neuroradiology, 2002. 23(5): p. 872–881.

45. Nossin-Manor, R., et al., Quantitative MRI in the very preterm brain: assessing tissue organization and myelination using magnetization transfer, diffusion tensor and T(1) imaging. Neuroimage, 2013. 64: p. 505–16.

46. Smyser, T.A., et al., Cortical Gray and Adjacent White Matter Demonstrate Synchronous Maturation in Very Preterm Infants. Cereb Cortex, 2016. 26(8): p. 3370–3378.

47. Ouyang, M., et al., Differential cortical microstructural maturation in the preterm human brain with diffusion kurtosis and tensor imaging. Proc Natl Acad Sci U S A, 2019. 116(10): p. 4681–4688.

48. Kliegman, R.M., et al., Epidemiologic study of necrotizing enterocolitis among low-birth-weight infants. Absence of identifiable risk factors. J Pediatr, 1982. 100(3): p. 440–4.

49. Kim, S.Y., et al., Disruption and Compensation of Sulcation-based Covariance Networks in Neonatal Brain Growth after Perinatal Injury. Cereb Cortex, 2020. 30(12): p. 6238–6253.

50. Kim, H., et al., Hindbrain regional growth in preterm newborns and its impairment in relation to brain injury. Hum Brain Mapp, 2016. 37(2): p. 678–88.

51. Kim, H., et al., NEOCIVET: Towards accurate morphometry of neonatal gyrification and clinical applications in preterm newborns. Neuroimage, 2016. 138: p. 28–42.

52. Liu, M., et al. A Skeleton and Deformation Based Model for Neonatal Pial Surface Reconstruction in Preterm Newborns. in 2019 IEEE 16th International Symposium on Biomedical Imaging (ISBI 2019). 2019. IEEE.

53. Smith, S.M., et al., Advances in functional and structural MR image analysis and implementation as FSL. Neuroimage, 2004. 23 Suppl 1: p. S208–19.

54. Wang, H. and P.A. Yushkevich, Multi-atlas segmentation with joint label fusion and corrective learning-an open source implementation. Front Neuroinform, 2013. 7: p. 27.

55. Smith, S.M., Fast robust automated brain extraction. Hum Brain Mapp, 2002. 17(3): p. 143–55.

56. Klein, A., et al., Evaluation of 14 nonlinear deformation algorithms applied to human brain MRI registration. Neuroimage, 2009. 46(3): p. 786–802.

57. Tzourio-Mazoyer, N., et al., Automated anatomical labeling of activations in SPM using a macroscopic anatomical parcellation of the MNI MRI single-subject brain. Neuroimage, 2002. 15(1): p. 273–89.

58. Dvornek, N.C., et al., Learning Generalizable Recurrent Neural Networks from Small Task-fMRI Datasets. Med Image Comput Comput Assist Interv, 2018. 11072: p. 329–337.

59. McKinstry, R.C., et al., Radial organization of developing preterm human cerebral cortex revealed by non-invasive water diffusion anisotropy MRI. Cereb Cortex, 2002. 12(12): p. 1237–43.

60. Righart, R., et al., Cortical pathology in multiple sclerosis detected by the T1/T2-weighted ratio from routine magnetic resonance imaging. Ann Neurol, 2017. 82(4): p. 519–529.

61. Arshad, M., J.A. Stanley, and N. Raz, Test-retest reliability and concurrent validity of in vivo myelin content indices: Myelin water fraction and calibrated T1 w/T2 w image ratio. Hum Brain Mapp, 2017. 38(4): p. 1780–1790.

62. Petracca, M., et al., Laminar analysis of the cortical T1/T2-weighted ratio at 7T. Neurol Neuroimmunol Neuroinflamm, 2020. 7(6).

63. Huppi, P.S., et al., Microstructural development of human newborn cerebral white matter assessed in vivo by diffusion tensor magnetic resonance imaging. Pediatr Res, 1998. 44(4): p. 584–90.

64. Back, S.A., et al., Arrested oligodendrocyte lineage progression during human cerebral white matter development: dissociation between the timing of progenitor differentiation and myelinogenesis. J Neuropathol Exp Neurol, 2002. 61(2): p. 197–211.

65. Drobyshevsky, A., et al., Developmental changes in diffusion anisotropy coincide with immature oligodendrocyte progression and maturation of compound action potential. J Neurosci, 2005. 25(25): p. 5988–97.

66. Wimberger, D.M., et al., Identification of “premyelination” by diffusion-weighted MRI. J Comput Assist Tomogr, 1995. 19(1): p. 28–33.

67. de Bruine, F.T., et al., Tractography of developing white matter of the internal capsule and corpus callosum in very preterm infants. Eur Radiol, 2011. 21(3): p. 538–47.

68. Bava, S., et al., Longitudinal characterization of white matter maturation during adolescence. Brain Res, 2010. 1327: p. 38–46.

69. Kersbergen, K.J., et al., Microstructural brain development between 30 and 40 weeks corrected age in a longitudinal cohort of extremely preterm infants. Neuroimage, 2014. 103: p. 214–224.

70. Miller, S.P., et al., Serial quantitative diffusion tensor MRI of the premature brain: development in newborns with and without injury. J Magn Reson Imaging, 2002. 16(6): p. 621–32.

71. Partridge, S.C., et al., Diffusion tensor imaging: serial quantitation of white matter tract maturity in premature newborns. Neuroimage, 2004. 22(3): p. 1302–14.

72. Oishi, K., et al., Multi-contrast human neonatal brain atlas: application to normal neonate development analysis. Neuroimage, 2011. 56(1): p. 8–20.

73. Dubois, J., et al., Assessment of the early organization and maturation of infants’ cerebral white matter fiber bundles: a feasibility study using quantitative diffusion tensor imaging and tractography. Neuroimage, 2006. 30(4): p. 1121–32.

74. Lobel, U., et al., Diffusion tensor imaging: the normal evolution of ADC, RA, FA, and eigenvalues studied in multiple anatomical regions of the brain. Neuroradiology, 2009. 51(4): p. 253–63.

75. Schmithorst, V.J. and W. Yuan, White matter development during adolescence as shown by diffusion MRI. Brain Cogn, 2010. 72(1): p. 16–25.

76. Volpe, J.J., Neurology of the newborn. Major Probl Clin Pediatr, 1981. 22: p. 1–648.

77. Huttenlocher, P.R. and A.S. Dabholkar, Regional differences in synaptogenesis in human cerebral cortex. J Comp Neurol, 1997. 387(2): p. 167–78.

78. Edde, M., et al., Functional brain connectivity changes across the human life span: From fetal development to old age. J Neurosci Res, 2021. 99(1): p. 236–262.

79. Canini, M., et al., Subcortico-Cortical Functional Connectivity in the Fetal Brain: A Cognitive Development Blueprint. Cerebral Cortex Communications, 2020. 1(1).

80. Wright, R., et al., Automatic quantification of normal cortical folding patterns from fetal brain MRI. Neuroimage, 2014. 91: p. 21–32.

81. Yu, Q., et al., Structural Development of Human Fetal and Preterm Brain Cortical Plate Based on Population-Averaged Templates. Cereb Cortex, 2016. 26(11): p. 4381–4391.

82. Deipolyi, A.R., et al., Comparing microstructural and macrostructural development of the cerebral cortex in premature newborns: diffusion tensor imaging versus cortical gyration. Neuroimage, 2005. 27(3): p. 579–86.

83. Ball, G., et al., Development of cortical microstructure in the preterm human brain. Proc Natl Acad Sci U S A, 2013. 110(23): p. 9541–6.

84. Kinney, H.C., et al., Myelination in the developing human brain: biochemical correlates. Neurochem Res, 1994. 19(8): p. 983–96.

85. Kinney, H.C., et al., Sequence of central nervous system myelination in human infancy. II. Patterns of myelination in autopsied infants. J Neuropathol Exp Neurol, 1988. 47(3): p. 217–34.

86. van den Heuvel, M.P., et al., The Neonatal Connectome During Preterm Brain Development. Cereb Cortex, 2015. 25(9): p. 3000–13.

87. Kawahara, J., et al., BrainNetCNN: Convolutional neural networks for brain networks; towards predicting neurodevelopment. Neuroimage, 2017. 146: p. 1038–1049.

88. Gui, L., et al., Longitudinal study of neonatal brain tissue volumes in preterm infants and their ability to predict neurodevelopmental outcome. Neuroimage, 2019. 185: p. 728–741.

89. McCrea, H.J. and L.R. Ment, The diagnosis, management, and postnatal prevention of intraventricular hemorrhage in the preterm neonate. Clin Perinatol, 2008. 35(4): p. 777-92, vii.

90. Anderson, A., et al., Modeling analysis of change in neurologic abnormalities in children born prematurely: a novel approach. J Child Neurol, 1999. 14(8): p. 502–8.

91. Arzoumanian, Y., et al., Diffusion tensor brain imaging findings at term-equivalent age may predict neurologic abnormalities in low birth weight preterm infants. AJNR Am J Neuroradiol, 2003. 24(8): p. 1646–53.

92. Albertine, K.H.J.C.i.p., Brain injury in chronically ventilated preterm neonates: collateral damage related to ventilation strategy. 2012. 39(3): p. 727–740.

93. Malavolti, A.M., et al., Bronchopulmonary dysplasia—impact of severity and timing of diagnosis on neurodevelopment of preterm infants: a retrospective cohort study. 2018. 2(1).

94. Van Marter, L.J., et al., Does bronchopulmonary dysplasia contribute to the occurrence of cerebral palsy among infants born before 28 weeks of gestation? 2011. 96(1): p. F20–F29.

95. Singer, L.T., et al., Preschool language outcomes of children with history of bronchopulmonary dysplasia and very low birth weight. 2001. 22(1): p. 19–26.

96. Natarajan, G., et al., Outcomes of extremely low birth weight infants with bronchopulmonary dysplasia: impact of the physiologic definition. 2012. 88(7): p. 509–515.

97. Short, E.J., et al., Cognitive and academic consequences of bronchopulmonary dysplasia and very low birth weight: 8-year-old outcomes. 2003. 112(5): p. e359–e359.

98. Singer, L., et al., A longitudinal study of developmental outcome of infants with bronchopulmonary dysplasia and very low birth weight. 1997. 100(6): p. 987–993.

99. Ouyang, M., et al., Delineation of early brain development from fetuses to infants with diffusion MRI and beyond. Neuroimage, 2019. 185: p. 836–850.

